# Resource sharing leads to the emergence of division of labour

**DOI:** 10.1101/2022.04.15.488476

**Authors:** Jan J. Kreider, Thijs Janzen, Abel Bernadou, Daniel Elsner, Boris H. Kramer, Franz J. Weissing

## Abstract

Division of labour occurs in a broad range of organisms. Yet, how division of labour can emerge in the absence of pre-existing interindividual differences is poorly understood. Using a simple but realistic model, we show that in a group of initially identical individuals, division of labour emerges spontaneously if returning foragers share part of their resources with other group members. In the absence of resource sharing, individuals follow an activity schedule of alternating between foraging and other tasks. If non-foraging individuals are fed by other individuals, their alternating activity schedule becomes interrupted, leading to task specialisation and the emergence of division of labour. Furthermore, nutritional differences between individuals reinforce division of labour. Such differences can be caused by increased metabolic rates during foraging or by dominance interactions during resource sharing. Our model proposes a plausible mechanism for the self-organised emergence of division of labour in animal groups of initially identical individuals. This mechanism could also play a role for the emergence of division of labour during the major evolutionary transitions to eusociality and multicellularity.

## Introduction

Division of labour – the non-random association of group-living individuals with tasks to be performed – is a pivotal aspect of social life in human and animal societies. Division of labour occurs in a broad range of organisms^1,2^. For instance, in eusocial insects, workers specialise in foraging, defending the nest, or nursing the brood^3^. In some species of birds, such as noisy miners, some helpers specialise in chick provisioning or mobbing nest predators^4^. In the Lake Tanganyika princess cichlid, helpers engage in predator defence, nest maintenance or egg care^5,6^. Within prides of lionesses, individuals perform different roles during hunting and territorial defence, and they can specialise to take care of the cubs^7^.

A central question for understanding social life is consequently how such division of labour can originate. The emergence of division of labour is typically modelled in response threshold models^8–14^. These models assume that individuals differ in thresholds that determine their likelihood to start performing a task when they perceive a task stimulus. Individuals with a low response threshold take on a task more readily, thereby decreasing the task stimulus; this prevents other individuals with a higher threshold from taking on the task as well. As a result, individuals specialise in those tasks for which they have a low threshold. However, task specialisation and division of labour only occur if there are pre-existing interindividual differences in response thresholds^15^. It remains poorly understood how self-organised division of labour can emerge within homogenous groups of highly similar or even identical individuals. Here, we present a model for the emergence of division of labour in animal groups of identical individuals. The model makes the realistic assumption that the nutrition level of individuals declines over time, and that a low nutrition level triggers foraging^16–22^ (Fig. 1a). In the absence of interindividual interactions, one would expect that individuals sequentially perform non-foraging behaviours (henceforth referred to as ‘nursing’) when nutrition levels are high and foraging behaviour when nutrition level is low. However, within groups of animals, individuals interact and resource sharing regularly occurs^23,24^. This could lead to a certain degree of specialisation on foraging and non-foraging behaviour, as foragers that give part of their resources away may soon have to forage again, while the receivers of the shared resources can delay foraging. We therefore considered various resource-sharing scenarios (Fig. 1b), in order to investigate whether this common mechanism is sufficient to generate a high degree of task specialisation and division of labour.

**Fig 1.**
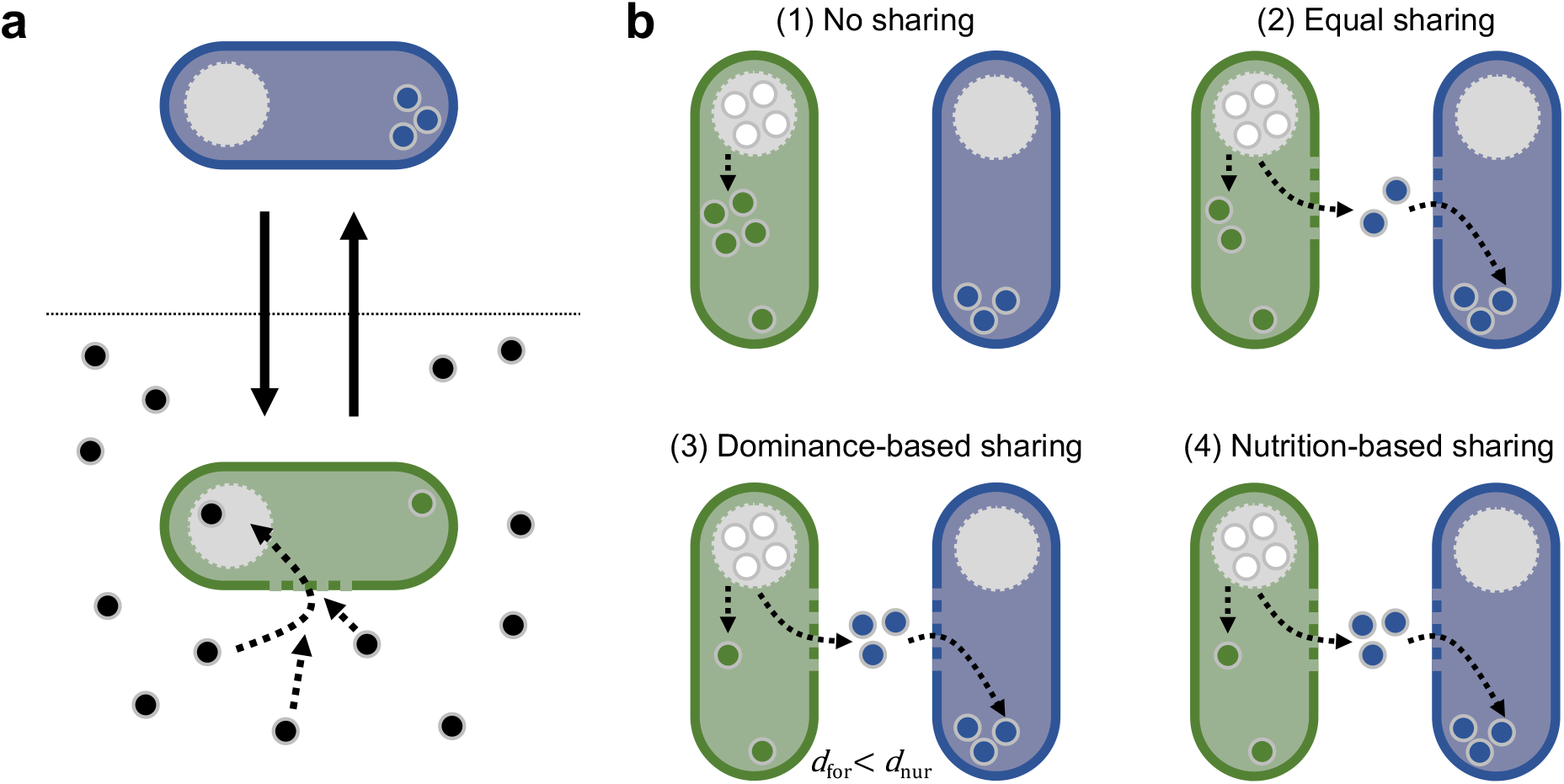
Task switching and four resource sharing scenarios. **(a)** Our model assumes that individuals can switch (solid arrows) between two states: foraging (green) and nursing (blue). While foraging, individuals retrieve resources (black dots) from the environment and store these resources in a temporary storage organ (grey circle). As long as resources are in the storage organ, they can in principle be shared with other individuals. Once they are integrated into the individual’s body (green dots in foraging individuals; blue dots in nursing individuals) they contribute to the individual’s nutrition level and can no longer be shared. As foraging is triggered by a low nutrition level, foraging individuals have on average lower nutrition levels than nursing individuals. **(b)** Four resource sharing scenarios considered in our study: (1) *No sharing*: Resources are not shared; foraging individuals consume all resources themselves. (2) *Equal sharing*: Foraging individuals share the collected resources equally between themselves and a nursing individual. (3) *Dominance-based sharing*: Each individual has a pre-assigned dominance value and resources are shared in relation to these dominance values: The dominant individual obtains a larger proportion of the collected resources. (4) *Nutrition-based sharing*: As in (3), but now the dominance level of an individual is not pre-assigned and constant, but proportional to the individual’s nutrition level.

## Results

### Resource sharing is sufficient for the emergence of division of labour

We ran 20 replicate simulations for each resource sharing scenario. We quantified the degree of division of labour that emerges in the simulations using the metric *D* that was introduced by Duarte and colleagues^25^ (see Methods for details and Fig. S1 in the Supplement for other division-of-labour metrics). *D* ranges from −1 to +1, where −1 indicates the strict alternation between tasks, 0 indicates random switching between tasks, and +1 indicates task specialisation^25^. Figure 2a shows that in the absence of resource sharing (Fig. 1b; (1) No sharing), the division-of-labour metric *D* takes on the extreme value −1, implying that individuals follow an activity schedule of alternating between tasks. Nursing individuals that start to forage to obtain resources keep all the resources for themselves. Consecutively, they nurse until their nutrient level has dropped to a threshold that induces them to start foraging again. As shown in Figure 2b, the outcome is very different when resources are shared between foraging and nursing individuals (Fig. 1b; (2) Equal sharing). Now the metric D reaches positive values above 0.5, indicating that individuals specialize on foraging or nursing for longer stretches of time.

**Fig 2.**
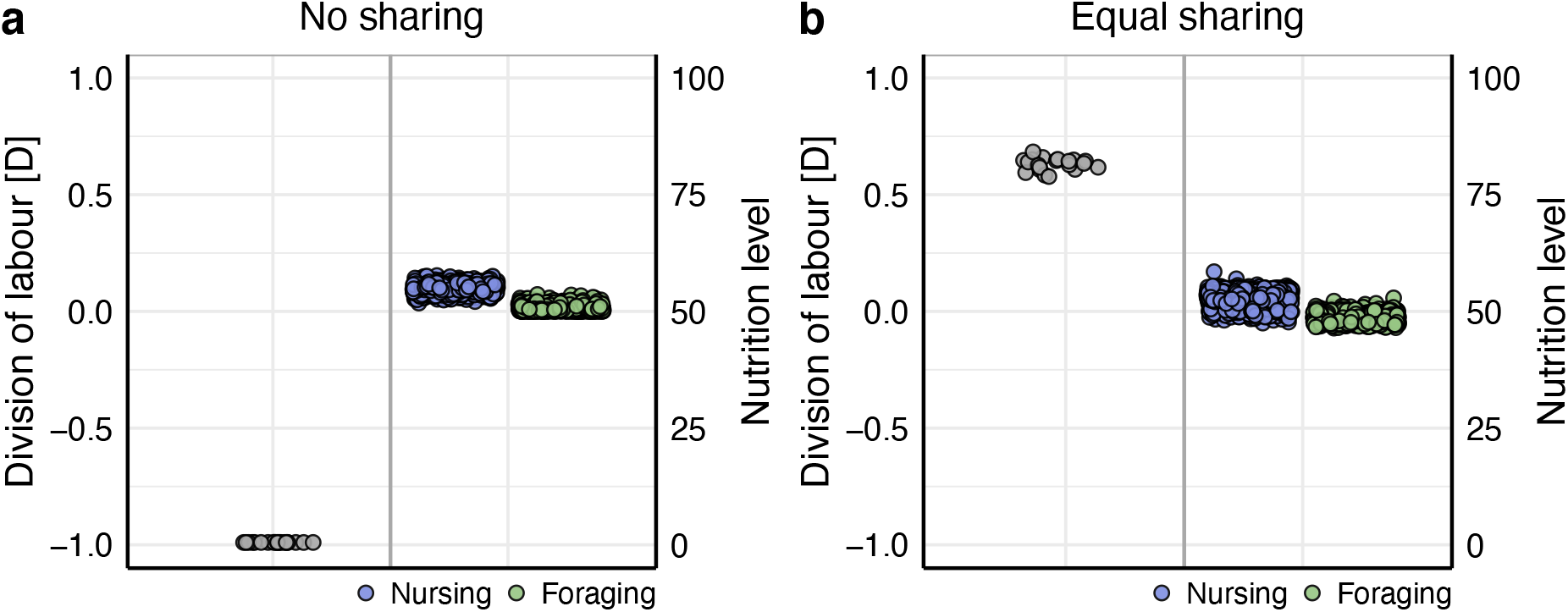
Division of labour and nutrition levels of nursing and foraging individuals under no resource sharing and equal sharing. **(a)** *No sharing*. In the absence of resource sharing, *D* = −1 which indicates that individuals alternate between nursing and foraging. As foraging is triggered by a decline in the nutrition level, nursing individuals have a higher nutrition level than foragers. **(b)** *Equal sharing*. If foraging individuals share their collected resources equally with nursing individuals, *D* ≈ 0.6 which represents an intermediate level of division of labour. Each grey dot (left panel) is the division of labour metric from a replicate simulation (*n* = 20). Blue (nursing) and green (foraging) dots (right panel) represent the nutrition levels of all individuals at the end of all replicate simulations.

### Division of labour is reinforced if foragers have a higher metabolic rate

As metabolic rates can differ with task^26,27^, we tested how such differences affect the emergence of division of labour. Even if the metabolic rate associated with nursing is only slightly lower (90% or 95%) than the metabolic rate associated with foraging, division of labour is strongly reinforced (Fig. 3a), leading to a bimodal distribution of nutrition levels (Fig. 3b). Conversely, if metabolic rates associated with nursing are higher compared to foraging, division of labour becomes weaker. Similarly, when the duration of nursing is shorter than the duration of foraging, division of labour is reinforced because this decreases the amount of energy that nursing individuals metabolise during task performance relative to foraging individuals (Fig. S2).

**Fig 3.**
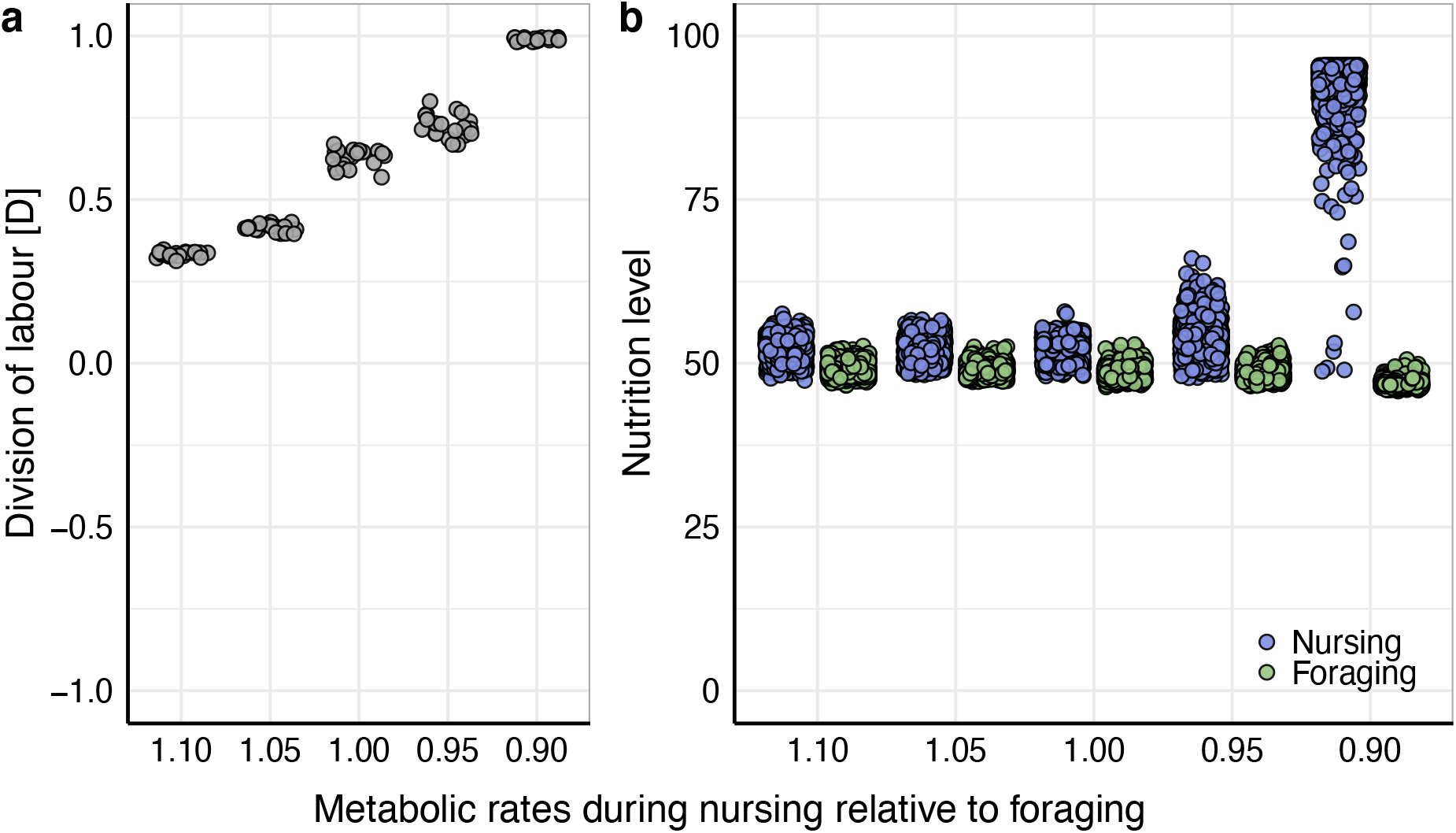
Division of labour when the metabolic costs of foraging and nursing differ. **(a)** If metabolic rates during nursing are lower relative to foraging, division of labour is reinforced. Conversely, division of labour is weaker when metabolic rates during nursing are higher than during foraging. **(b)** As their metabolic rates become lower, the nutrition level of nursing individuals deviates more strongly from those of foraging individuals. For graphical conventions, see Fig. 2.

### Dominance relationships reinforce division of labour, especially when dominance is related to nutritional status

Resource sharing is often not equal but associated with dominance^18,28–30^. In order to investigate uneven resource sharing through dominance effects, we assigned constant dominance values to the individuals in the simulation at initialisation (Fig. 1b; (3) Dominance-based sharing). Under such dominance-based differences in resource intake, division of labour reaches maximal levels and nutrition levels of foraging and nursing individuals are bimodally distributed (Fig. 4a1). As individuals with larger dominance values obtain more resources during sharing, they rapidly are fixed into nursing whereas those with low dominance values consistently forage. Individuals with intermediate dominance values either forage or nurse (Fig. 4a2), which is stochastically driven by the random interactions that those individuals have at the start of a simulation. The model scenario considered in Fig. 4a has the drawback that division of labour is based on pre-existing differences (in dominance) between individuals. However, the same result can also be obtained in a population of initially identical individuals, if dominance is caused by differences in nutrition level (Fig. 1b; (4) Nutrition-based sharing). As shown in Figure 4b1, division of labour again reaches maximal levels, and the nutrition levels of foraging and nursing individuals are bimodally distributed. At the start of the simulation, nutrition levels of individuals diverge relatively slowly but once small differences are present, a full divergence of nutrition levels is rapidly achieved (Fig. 4b2). In Figure S3, we show that low levels of division of labour even emerge if individuals with low nutrition levels obtain more resources during sharing. This represents a more altruistic sharing scenario.

**Fig 4.**
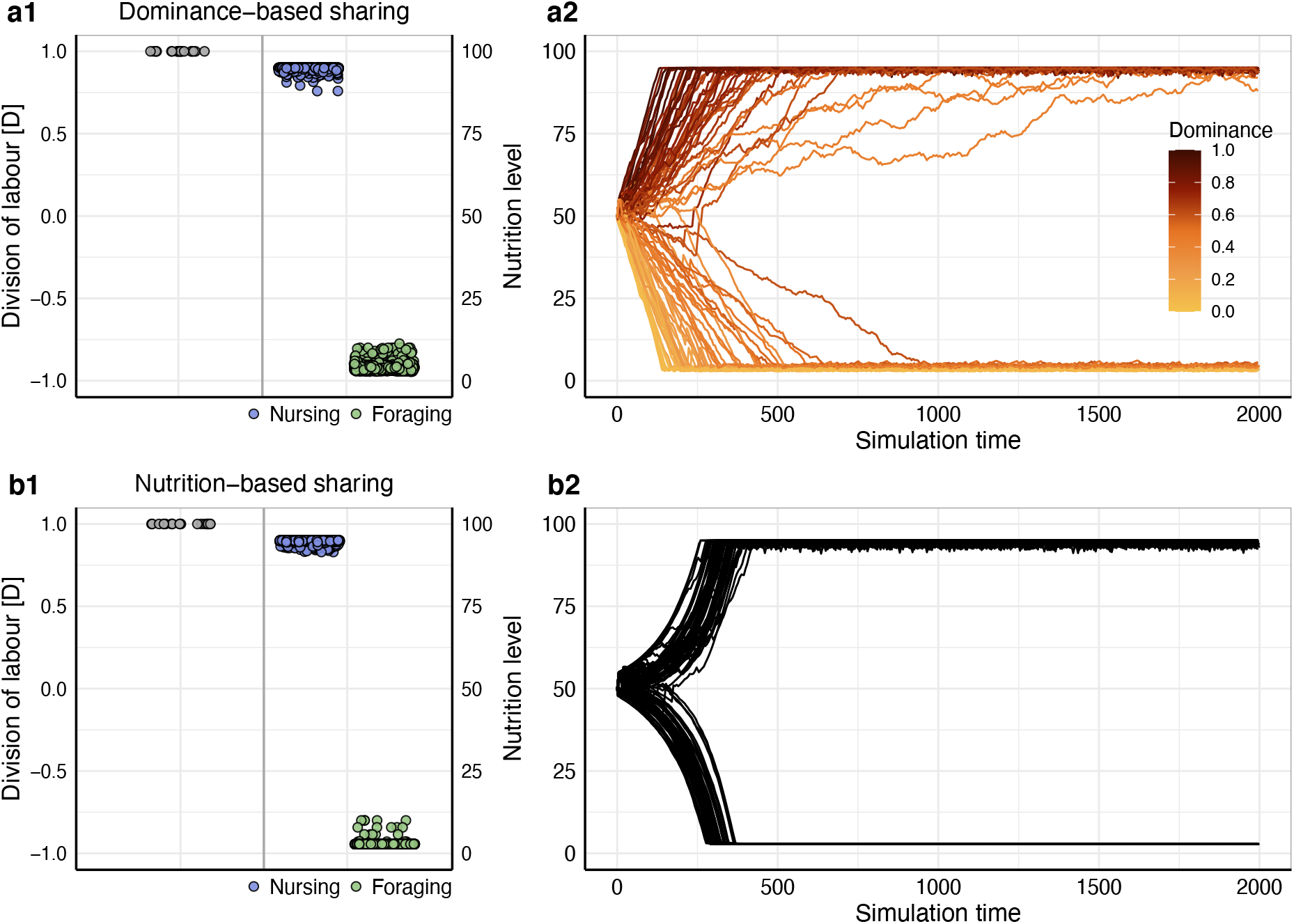
Division of labour, nutrition level and dominance of nursing and foraging individuals. **(a1)** *Dominance-based sharing*. Division of labour reaches maximal levels of *D* = 1. Nursing and foraging individuals exhibit a bimodal distribution of nutrition levels. **(a2)** Nutrition levels of individuals diverge over simulation time, depending on the individual’s dominance. **(b1)** *Nutrition-based sharing*. Division of labour again reaches maximal levels of *D* = 1, and nutrition levels exhibit a bimodal distribution. **(b2)** Nutrition levels of individuals diverge over simulation time. **(a1 + b1)** For graphical conventions, see Fig. 2. **(a2 + b2)** Each line shows the nutrition level of an individual over the first 20% of simulation time from a representative replicate simulation, coloured according to the individual’s dominance value if dominance values were pre-assigned.

### Division of labour emerges irrespective of group size

Specialisation has been suggested to increase with group size^11,31,32^ but this is not supported unequivocally^25,33^. We therefore altered group size to investigate its effect on the emergence of division of labour. Figure 5 shows that division of labour emerges independently of group size. However, in small groups, the degree of division of labour may strongly depend on details (such as whether group size is even or odd). Fig. S4 further elaborates on this.

**Fig 5.**
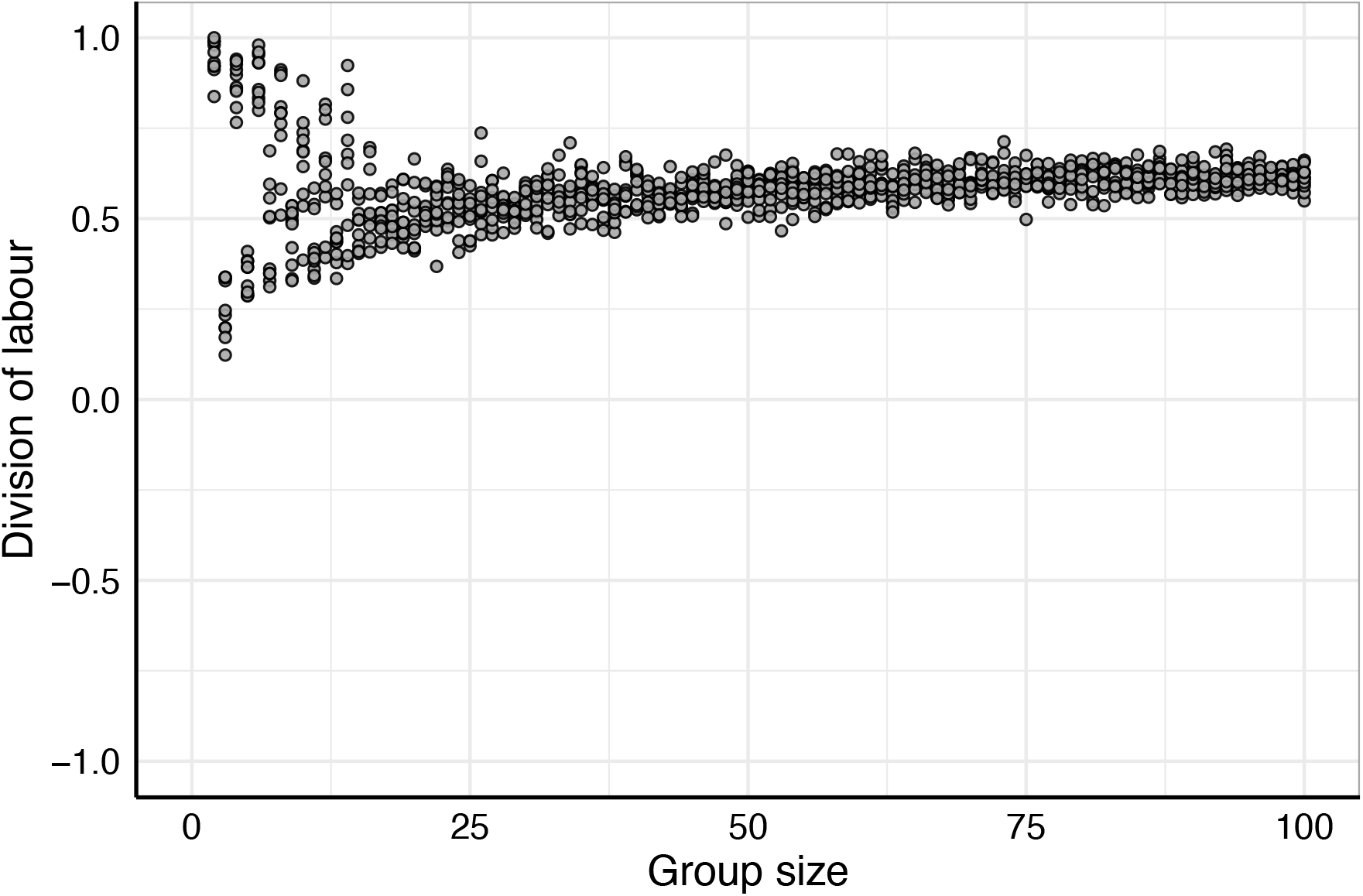
Effect of group size on division of labour in the equal sharing scenario. Division of labour emerges irrespective of group size. Each dot shows the division of labour metric from a replicate simulation per group size (*n* = 10).

## Discussion

The emergence of division of labour is typically modelled in response threshold models^8–14^. Here, we demonstrate that resource sharing is an alternative mechanism leading to the emergence of division of labour. Our model reveals that even in a group of initially identical individuals, resource sharing can result in very strong task specialisation and division of labour. Pre-existing differences between individuals can enhance division of labour, but they are not required for its emergence. In contrast, most response threshold models are based on the assumption that, from the start, individuals differ in their threshold values.

Our model is based on two general and plausible assumptions. First, individual nutrient levels decline in the course of time, inducing individuals to go out foraging once they become very low. In the absence of interactions with other individuals, this results in an activity schedule where individuals alternate between nursing (when nutrient levels are declining) and foraging (when nutrient levels are replenished). If, second, foragers share some of their collected resources with other individuals, this activity schedule becomes interrupted due to two feedbacks: nursing individuals may be fed before perceiving a hunger signal, thus retarding their next foraging bout; and foraging individuals cannot use the shared resources to fully replenish their nutrient level, thus reducing the time until the hunger signal induces them to start foraging again. These feedbacks lead to task specialisation and division of labour, because nursing individuals that are fed by others are likely to continue nursing whereas foraging individuals, who feed others, are likely to proceed with foraging.

Several studies have proposed that division of labour in social insects are associated with nutritional differences between foragers and nurses^16–22,34–36^. In our model, (a) division of labour leads to a situation where nutrient levels of nurses are higher than those of foragers, and (b) that division of labour is enhanced when the metabolic rate while foraging is higher than the metabolic rate while nursing. However, in our model pre-existing nutritional differences are not required for the emergence of division of labour – the crucial ingredient is resource sharing. In social insects, resources can, for instance, be transferred mouth-to-mouth via trophallactic exchanges^24^. However, for the emergence of division of labour it is not required that resources are shared altruistically. Foragers could, for instance, deposit part of the resources in the nest (destined to be fed to the brood at a later stage), where they are consumed by other individuals instead.

Our model highlights the importance of dominance interactions and task-specific differences in nutrition level for the emergence of strong non-reproductive division of labour. Dominance and nutrition level also play an important role in the determination of reproductive abilities in social insects^28–30,37,38^ and in social spiders^39^, and thus in reproductive division of labour. For instance, in paper wasps, breeder and helper roles are likely determined by nutrition^40^, as also indicated by the differential expression of storage proteins in breeders and helpers^41,42^. Furthermore, in eusocial insects, caste is often determined by a nutrition-dependent developmental switch^43^, and across eusocial insects, castes differentially express genes coding for the storage protein vitellogenin^36,44–47^. Thus, differences in nutrition level that emerge in our model due to dominance interactions during resource sharing could also play an important role for the evolution of eusociality.

Response threshold models and empirical studies^11,31,32^ (though not all^25,33^) suggest that task specialisation and division of labour are most pronounced in large groups. In our model, division of labour emerges even in very small groups. In fact, division of labour can be stronger in very small groups (e.g. groups of two) than in larger groups. Equivalently, experiments have shown that division of labour emerges in paired associations of ants^18,48^, and the sexes of some species of birds exhibit strong division of labour between the breeding female and the foraging male^49–51^. Consequently, the strength of division of labour does not necessarily depend on group size alone but also the details of social interactions, such as the number of individuals with which foragers share resources, are crucial to be considered.

Lastly, resource sharing could be a mechanism for the emergence of division of labour beyond animal groups. For instance, in some green algae, cells are specialised on obtaining different resources – carbon or nitrogen – for the colony^52^. Resource sharing could thus play a role in the emergence and regulation of cellular division of labour during the major evolutionary transition to multicellularity^53,54^.

Overall, our model demonstrates that self-organised division of labour emerges through resource sharing between identical individuals. Given the omnipresence of resource sharing in biological systems – for instance, in group-living animals, eusocial insects, human hunter-gatherer societies, or within multicellular organisms – our model suggests a mechanism that could explain the emergence of division of labour across a broad range of organisms.

## Methods

### The model

We developed an individual-based simulation model in continuous time. Each simulation represents a group of *N* individuals (*N* = 100, unless stated otherwise). Individuals either forage to obtain resources or perform other tasks to which we refer as ‘nursing’. Each simulation had a duration of *T* time steps (*T* = 10 000).

### Nutrition level of individuals

Individuals possess an internal state variable *n* that reflects their nutrition level and ranges from 0 to *n*_max_ (*n*_max_ = 100). At the start of a simulation, all individuals are initialised with an identical nutrition level of *n*_init_ (*n*_init_ = 50). Thus, simulations start with identical individuals. Over time, the nutrition level of individuals decreases by a metabolic rate *m*_for_ when foraging, and *m*_nur_ when nursing (*m*_for_ = 1.0, *m*_nur_ = 1.0, unless stated otherwise).

### Task choice

An individual’s choice to forage or nurse depends on the individual’s nutrition level. We assume that individuals become more likely to forage as their nutrition level declines. On average, individuals start foraging when their nutrition level reaches the critical value *μ*. However, we assume that individuals are not perfectly accurate in assessing their nutrition level. Consequently, we sample the exact nutrition level at which individuals start to forage from a normal distribution with mean *μ* and standard deviation *σ* (*μ* = 5 0, *σ* = 1). The probability to forage is checked upon completion of a task; foraging individuals do so after returning from a foraging trip, which has a fixed length of *t*_for_ (*t*_for_ = 5); similarly, nursing individuals do so after finishing their task, which has a fixed length of *t*_nur_ (*t*_nur_ = 5), and after obtaining resources from foragers.

### Nutrition transfers

Foragers obtain *R* resources while foraging (*R* = 10). We implemented four different model scenarios of resource sharing. (1) *No sharing*: Upon return from a foraging trip, foraging individuals keep all of the resources they obtained. (2) *Equal sharing*: Upon return from a foraging trip, foraging individuals evenly share resources with *i* nursing individuals. The number of resources that the individuals engaged in the interaction obtain is thus *R*/(1 + i) because one foraging individuals interacts with *i* nursing individuals. We assume that *i* = 1 but relax this assumption in the supplementary materials (Fig. S4). (3) *Dominance-based sharing*: Individuals are assigned a dominance value upon initialisation of the simulation, randomly sampled from a uniform distribution ranging from 0 to 1. The number of resources obtained by the individuals during the interaction is dependent on their dominance value, such that more dominant individuals obtain more resources than less dominant individuals. The number of resources obtained by individual *k* is calculated using the softmax function

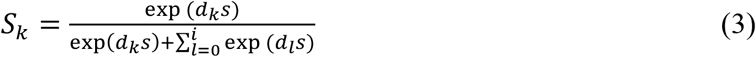

where *d_k_* is the dominance value of focal individual *k, d_I_* is the dominance value of the other *i* individuals and *s* is a parameter for the softmax function. If *s* = 0, the model reduces to the equal sharing scenario, if *s* < 0, individuals with the lowest dominance values receive a disproportionate number of resources, and if *s* > 0, individuals with the highest dominance values receive a disproportionate number of resources. For the simulations with a dominance effect, we used *s* = 1, which results in the most dominant individual getting slightly more than their expected share based on their relative dominance. (4) *Nutrition-based sharing*: The dominance status of an individual is given by its nutrition level, relative to the maximum nutrition level possible, i.e. *d_k_* is now given by *n/n*_max_. Otherwise, interactions proceed as in scenario 3, using equation (3) to determine resource distribution among individuals. Again, we used *s* = 1 (for simulations with other values of *s*, see Fig S3). As dominance again determines resource distribution among individuals, individuals who have a higher nutrition level obtain more resources than individuals with a lower nutrition level.

When a foraging individual returns from a foraging trip, but no nursing individuals who can obtain resources are available, the foraging individual consumes all of the resources itself, independently of the sharing scenario.

### Quantifying division of labour

We quantified the degree of division of labour that emerges in the simulations using the division of labour metric from Duarte et al.^25^, which is calculated as follows. 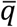 is the average probability of individuals in the group to perform identical tasks in two subsequent time steps. To obtain 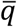, first, the probability of performing identical tasks in two subsequent time steps is calculated for each individual, and then averaged over all individuals in the group. *p*_1_ and *p*_2_ are the proportions of time that individuals in the group spend on task 1 (e.g. foraging) and task 2 (e.g. nursing). Thus, 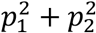 is the probability that individuals randomly consecutively perform the same tasks. Division of labour is then

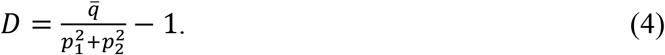

Here, −1 is subtracted to map the metric into the range of −1 to +1. A result of −1 indicates alternation between tasks, 0 indicates random switching and +1 indicates task specialisation^25^. Results based on two other division of labour metrics by Gautrais et al.^31^ and Gorelick et al.^55^ can be found in the supplementary materials (Fig. S1). We calculated division of labour metrics over the last 10% of simulation time to avoid measuring initialisation effects.

### Model analysis

The model was implemented in C++ and compiled with g++ 9.3.0. Model results were analysed and visualised in R 4.1.0^56^ using the packages *ggplot2*^57^, *gridExtra*^58^, *cowplot*^59^ and *MetBrewer*^60^.

## Acknowledgements

We thank Hanno Hildenbrandt for helping us debug the model and increase simulation efficiency. We thank the Center for Information Technology of the University of Groningen for providing access to the Peregrine high performance computing cluster. JJK was supported by an Adaptive Life grant by the University of Groningen. FJW acknowledges funding from the European Research Council (ERC Advanced Grant No. 789240).

## Author contributions

Conceptualisation: JJK, TJ, AB, DE, BHK, FJW; Implementation: TJ, JJK; Model analysis: JJK, DE; Writing: JJK, TJ, AB, DE, BHK, FJW.

