## Supplement for "Resource sharing leads to the emergence of division of labour"

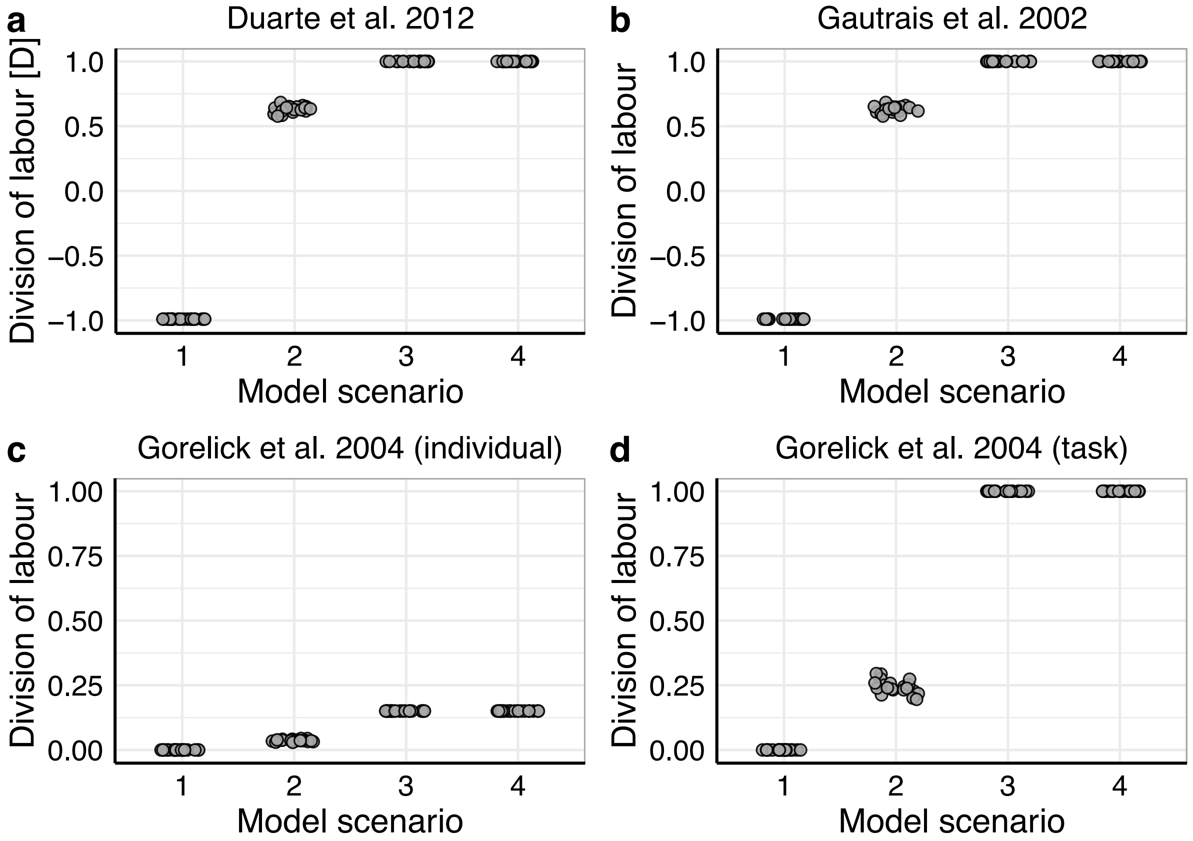


**Fig S1 | Comparison of different division of labour metrics in the four model scenarios.** **(a)** The division of labour metric of Duarte et al.[19] (used in the main manuscript), and **(b)** the division of labour metric of Gautrais et al.[53] are measures of individual specialisation and returns values of +1 when no task switches occur and -1 when tasks are performed in alternation. **(c + d)** The division of labour metrics from Gorelick et al.[54] are based on information theory and show **(c)** the predictability of the individual given that the task is known and **(d)** the predictability of the task given that the individual is known. High predictabilities are indicated by values close to 1 and low predictabilities by values close to 0. In **(c)**, the predictability of the individual when knowing the task always remains low because there are only two tasks but 100 individuals. Each dot shows the division of labour metric from a replicate simulation per model scenario ($n = 20$).

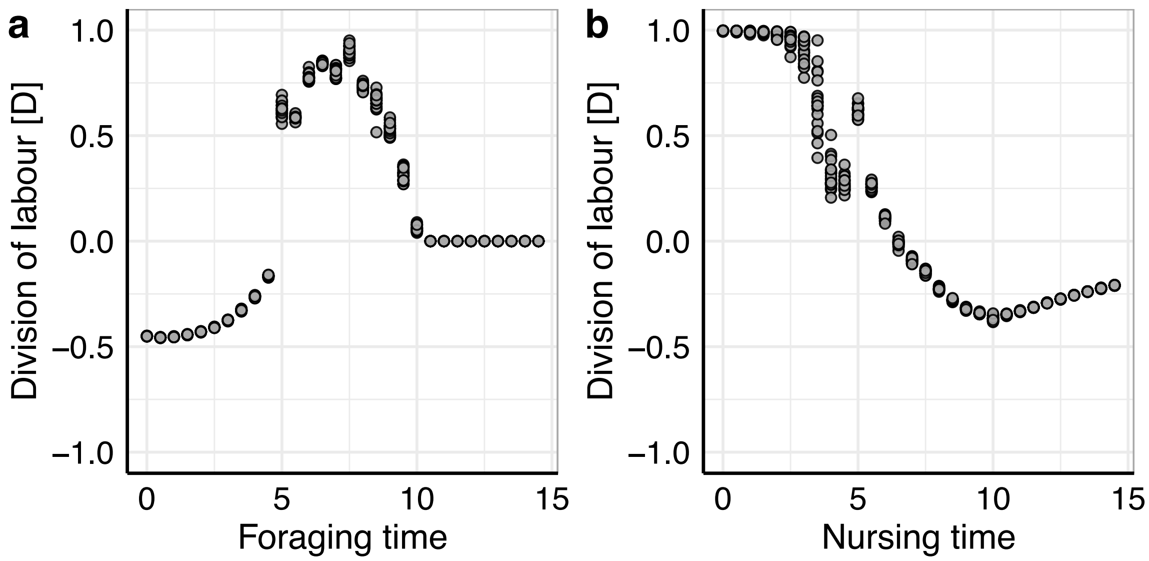


**Fig S2 | The effect of foraging and nursing time on the emergence of division of labour in the equal sharing scenario.** All simulations in the main manuscript were run with foraging and nursing times of 5. **(a)** If foraging time is lower, division of labour decreases (to values below zero) because foraging individuals metabolise fewer resources during foraging than they obtain on a foraging trip, thus leading to a resource accumulation in foraging individuals. If foraging time is higher, division of labour increases because foraging individuals metabolise more resources during foraging than they obtain. The division-of-labour metric becomes 0 when foraging time exceeds 10 because foraging individuals metabolise all of the resources during foraging that they obtain, even if they do not share resources with others; thus, all individuals are foraging. **(b)** If nursing time is higher, nursing individuals metabolise more resources during nursing than they obtain during sharing and thus they start foraging after nursing, leading to decreased division of labour. Each dot shows the division of labour metric from a replicate simulation per foraging or nursing time ($n = 20$).

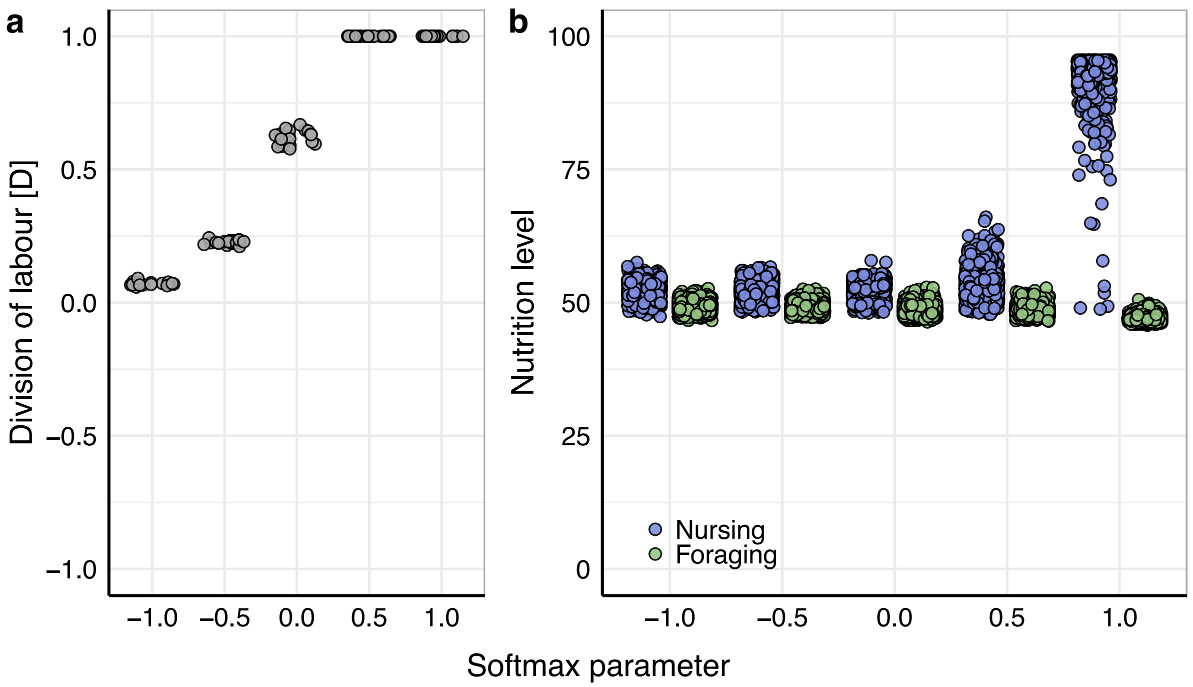


**Fig S3 | Division of labour for different values of the softmax parameter *s* in the nutrition-based sharing scenario. (a)** If individuals with higher nutrition level obtain more resources during sharing ($s > 0$), a high degree of division of labour emerges. However, even if individuals with lower nutrition level obtain more resources during sharing ($s < 0$), division of labour is still (weakly) positive. **(b)** If individuals with higher nutrition level obtain more resources during sharing ($s > 0$) the nutrition levels of foraging and nursing individuals are bimodally distributed; this is not the case for $s \leq 0$. Each grey dot (left panel) is the division of labour metric from a replicate simulation ($n = 20$). Blue (nursing) and green (foraging) dots (right panel) represent the nutrition levels of all individuals at the end of all replicate simulations.

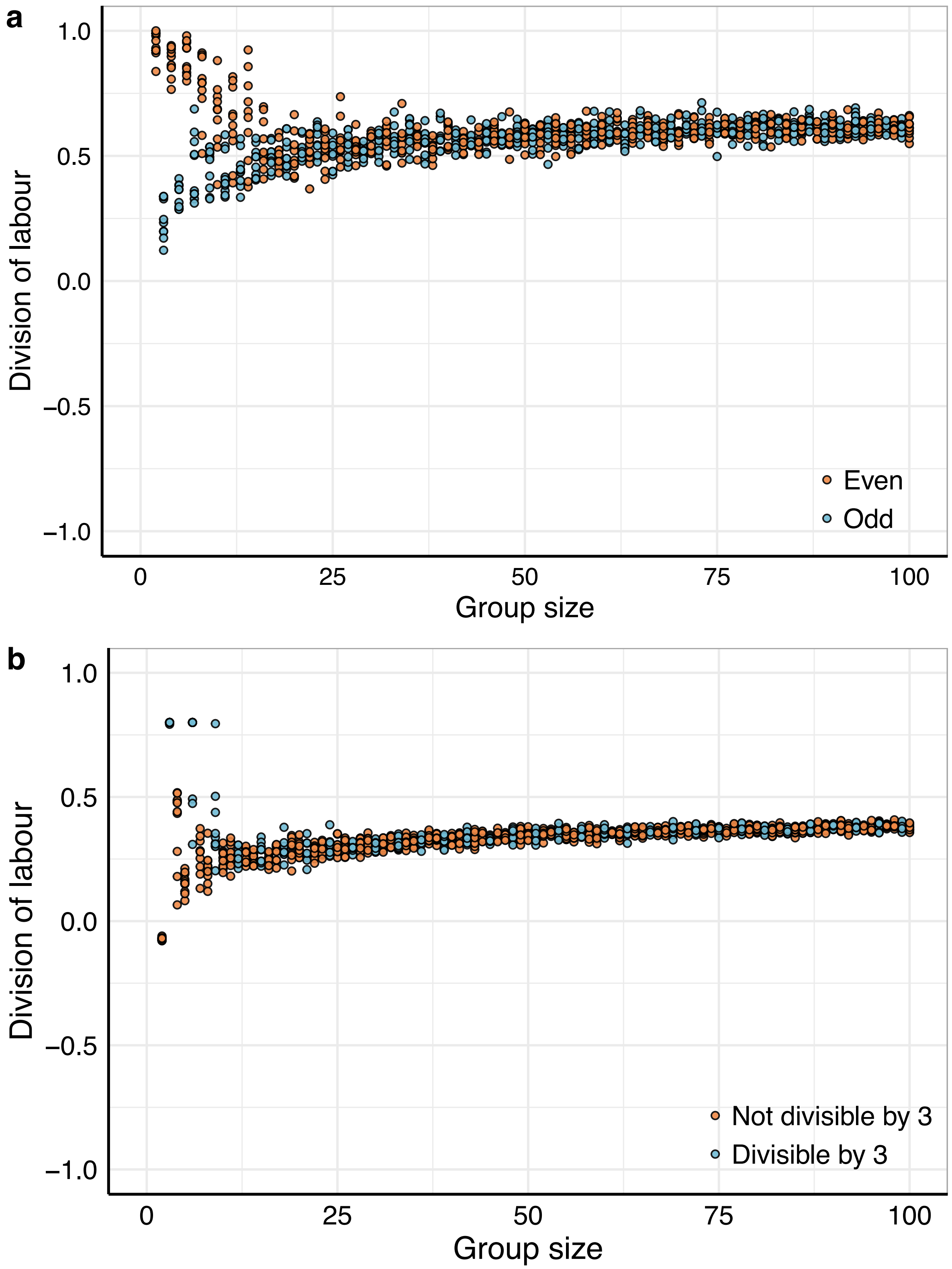


**Fig S4 | Division of labour as an effect of group size in the equal sharing. (a)** Division of labour is stronger if groups consist of even numbers of individuals compared to odd numbers of individuals. In groups of odd sizes, one individual cannot obtain resources through resource sharing and consequently alternates between foraging and nursing. In small groups, this has a larger effect on division of labour than in larger groups. Consequently, division of labour is not necessarily weaker in small groups, but the interplay of group size and the number of interactions of individuals determines whether division of labour is weak or strong in small groups. **(b)** In these simulations, foraging individuals share resource with two nursing individuals instead of with just one. Again, the degree of division of labour at small group sizes is dependent on whether group size is divisible by $1 + i$ (the number of nursing individuals a foraging individual interacts with). In order to maintain the assumption of net resource intake equalling net resource consumption, we increased the resource amount from 10 to 15 because now a single foraging individual has to feed itself plus two nursing individuals. Each dot shows the division of labour metric from a replicate simulation per group size ($n = 10$).
